# Toxic metal uptake by oyster mushrooms grown in sugarcane bagasse

**DOI:** 10.1101/2023.02.07.523865

**Authors:** Tirthankar Saha, Sagnik Das, Snigdha Sau, Debarpita Datta, Sourima Kundu, Subham Saha, Shreya Chakraborty, Arup Kumar Mitra

**Affiliations:** St. Xavier’s College, Kolkata

**Keywords:** Fungiculture, Nutritional value, Oyster Mushrooms, Bagasse, Heavy metal ions, Biomagnification

## Abstract

Fungiculture or Mushroom Cultivation is rapidly expanding throughout the world. It is further catalyzed by its high nutritional value and increased market demand. Additionally, the method of cultivation does not require much space and the per unit production is very high compared to other crops. Bagasse is the byproduct left after extracting the juice from sugarcane. It has been speculated that bagasse can be an excellent substrate due to its high carbohydrate and mineral content (Hoa et al., 2015). In the experiment spawns of Oyster Mushroom were inoculated in bagasse collected from industries. Remarkable growth of mushrooms in short period of time was observed. However, the metal toxicity being our primary concern in this experiment, AAS (Atomic Absorption Spectrometry) Test was used to detect the heavy metal ion concentrations in both the bagasse and the fruiting bodies of mushrooms. The concentrations of metals calculated are as follows, in bagasse (mg/kg): Cadmium - 0.015, Arsenic - 8.56, Copper-12.47, Chromium-9.17, Mercury - less than 0.01, and in mushroom (mg/kg): Cd - 0.022, As - 10.12, Cu - 14.29, Cr - 7.59, Hg - less than 0.01. These results indicated biomagnification of metal ion concentration in the mushrooms up taken from bagasse. These metals, in such high concentrations are lethal and causes many diseases in humans and other animals. Hence, from this experiment it can be deduced, that though bagasse is an excellent substrate for mushroom cultivation, the toxicity of metal ions present in it in such high concentrations overshadow the earlier benefits of bagasse as a good substrate. However, it has been proposed that if such metals are removed or their concentrations are lowered then the earlier mentioned benefits of bagasse are restored.

## 1. Introduction

Mushroom cultivation is on the rise since it can not only meet the dietary requirements, but also add to the income, especially for farmers with insufficient lands. Mushroom farming requires very little investment but reaps out high profits by more than 100% of the original investment.

The most fascinating part of it is that a small quantity of spawn when planted in suitable growth media can grow into highly profitable crop within almost 6 weeks even in a room, provided it has a damp and shaded environment. Mushrooms have also taken a huge leap to acquire a quite lucrative position in the modern culinary cuisine than any other food crops. There are many types of edible mushrooms suitable for cultivation in India, like - Button Mushroom (*Agaricus bisporus)*, Straw Mushroom (*Volvariella volvacea)*, Oyster Mushroom (*Pleurotus ostreatus*), etc. (Borah et al., 2020)

Our study here is mainly concentrated on oyster mushrooms which is a mushroom popularly grown in the South-East Asian countries. The conventional substrate for mushroom cultivation in these countries is straw but over the last two decades sugarcane and sorghum bagasse have emerged as one of the prominent and superior alternatives over straw. The quantity of mushroom that can be produced using the sugarcane bagasse is significantly higher than that of conventional straw, given the same quantity and condition (Hoa et al., 2015). At the same time the nutritional quality of the mushrooms grown on sugarcane bagasse is much higher than that of straw, which predominantly due to the presence of higher mineral and sugar content in sugarcane bagasse (Yadav et al., 2015; Agnihotri et al., 2010). Being an agricultural product bagasse has been found to contain metal ions and the mushroom being an edible product has to be ensured of possible toxic effect.

### 1.1. Metal ion toxicity

Heavy metals occur naturally in ecosystem with large variations in concentrations. In modern times anthropogenic sources of heavy metals, i.e., pollution have been introduced into the ecosystem. Pesticides and insecticides are most prone to contain heavy metals, thus presence of heavy metals in the substrate for food crop growth is of constitutional importance as once these food crops suffering from heavy metal contamination are up taken, it results in entry of these heavy metals in to the food chain and may also increase many folds by biomagnification (Demirbaş, 2002).

Hence the metal ions can cause severe effects in human lives leaving behind long term effects if not taken care of properly in due time.

In our in-vivo experimental study, the metal ion uptake by the common staple food crop *Pleurotus ostreatus* was studied and thus inferred to detect the entry of heavy metals into the food chain.

## 2. Method of cultivation of Oyster mushrooms in bagasse

Bagasse is collected after sugarcane is milled for juice extraction, which is mainly fibrous in nature and contains 60-80 % carbohydrates, primarily made up of lignin and cellulose (Mahmud & Anannya, 2021). The bagasse was boiled for 10 minutes (Anwar, 2010) so that, (i) the lignin and cellulose was softened to allow easy growth of mushroom, (ii) the bagasse was partially sterile, thereby inhibiting growth of other microbes and (iii) to allow tight packaging (Rivera, 2015).

The excess moisture was removed by heat drying followed by a 7-day lyophilization. The lyophilized bagasse was then packed and the moisture content was replenished to 40% (by weight) by adding sterile water.

A cellophane bag is used and successive 2-inch-thick layers of bagasse and spawns were spread out on it, this process was repeated till the layers were 8-10 inches thick.

The bag was twisted and sealed using strings to allow partial exchange of gases. A few (7-10) holes were made using a sharp tool to allow passage of air and the packets were kept in a dark area. Water was sprayed on all the slits everyday twice.

After 7 days white areas were observed, indicating that the spores have started germinating. After almost 10 days when all the spores had germinated, slits of 3-4 inches were made all over the side of the packet from where the mushrooms were expected to emerge (Moda et al., 2005).

## 3. Methods for metal ions toxicity test

The mushroom fruit bodies and the bagasse were dried up and converted to dust and were tested for presence of metal ions by Atomic Absorption Spectrometry (AAS). (Chirayil et al., 2017)

Each dust sample was weighed as 1 g for every metal to be tested and turned to ash. Then the ash was dissolved in 1 ml concentrated HNO_3_, evaporated to dryness, heated again at 400°C for 4 hours, treated with 1 ml concentrated H_2_SO_4_, 1 ml HNO_3_ and 1 ml H_2_O_2_ and then diluted with double deionized water up to a volume of 10 ml. Three blank samples were treated in the same way (Dospatliev, 2017).

In order to analyze a sample for its atomic constituents, the samples were volatilized by atomization; then the atoms were irradiated by optical radiation, and the radiation source could be an element-specific line radiation source or a continuum radiation source. Now, in order to separate the element-specific radiation from any other radiation emitted by the radiation source the radiation is passed through a monochromator, and it is finally measured by a detector (Ahmed, 2012). The standard addition procedure was used in all determination.

## 4. Results

### 4.1. Results of cultivation

After 17 days the whole packet turned white as fungal mycelium developed and covered the entire substrate (this is called spawn running stage, 25-30° C is optimum for this) and the mushrooms started bulging out from the slits. After 25 days it was seen that the pearl oyster mushrooms of varying sizes of fruit bodies grew from the slits. The mushroom was harvested by twisting the base of the mushroom. It could be harvested 3 times from each packet.

Total weight of fruiting bodies = 10.516 g.

### 4.2. Results for metal ions uptake

The mushroom and bagasse samples were tested for the heavy metals _ Cadmium, Arsenic, Copper, Chromium, Mercury. The obtained data is given in Table I.

**TABLE I.** Comparative study of concentration of heavy metal ions in bagasse and mushrooms grown in it. (Mean ± Standard Deviation) (n = 3)

| **TABLE I.** Comparative study of concentration of heavy metal ions in bagasse and mushrooms grown in it. (Mean ± Standard Deviation) (n = 3) | | | |
| --- | --- | --- | --- |
| Heavy metal | Concentration in bagasse (mg/kg, dry weight) | Concentration in mushroom (mg/kg, dry weight) | Bioaccumulation Factor |
| Cadmium (Cd) | 0.0153 ± 0.002 | 0.0227 ± 0.004 | 1.48 |
| Arsenic (As) | 8.2733 ± 0.594 | 10.0933 ± 0.074 | 1.22 |
| Copper (Cu) | 12.103 ± 0.661 | 14.0067 ± 0.535 | 1.16 |
| Chromium (Cr) | 9.0133 ± 0.289 | 7.6467 ± 0.211 | 0.85 |
| Mercury (Hg) | <0.01 | <0.01 | - |

**Figure 1:**
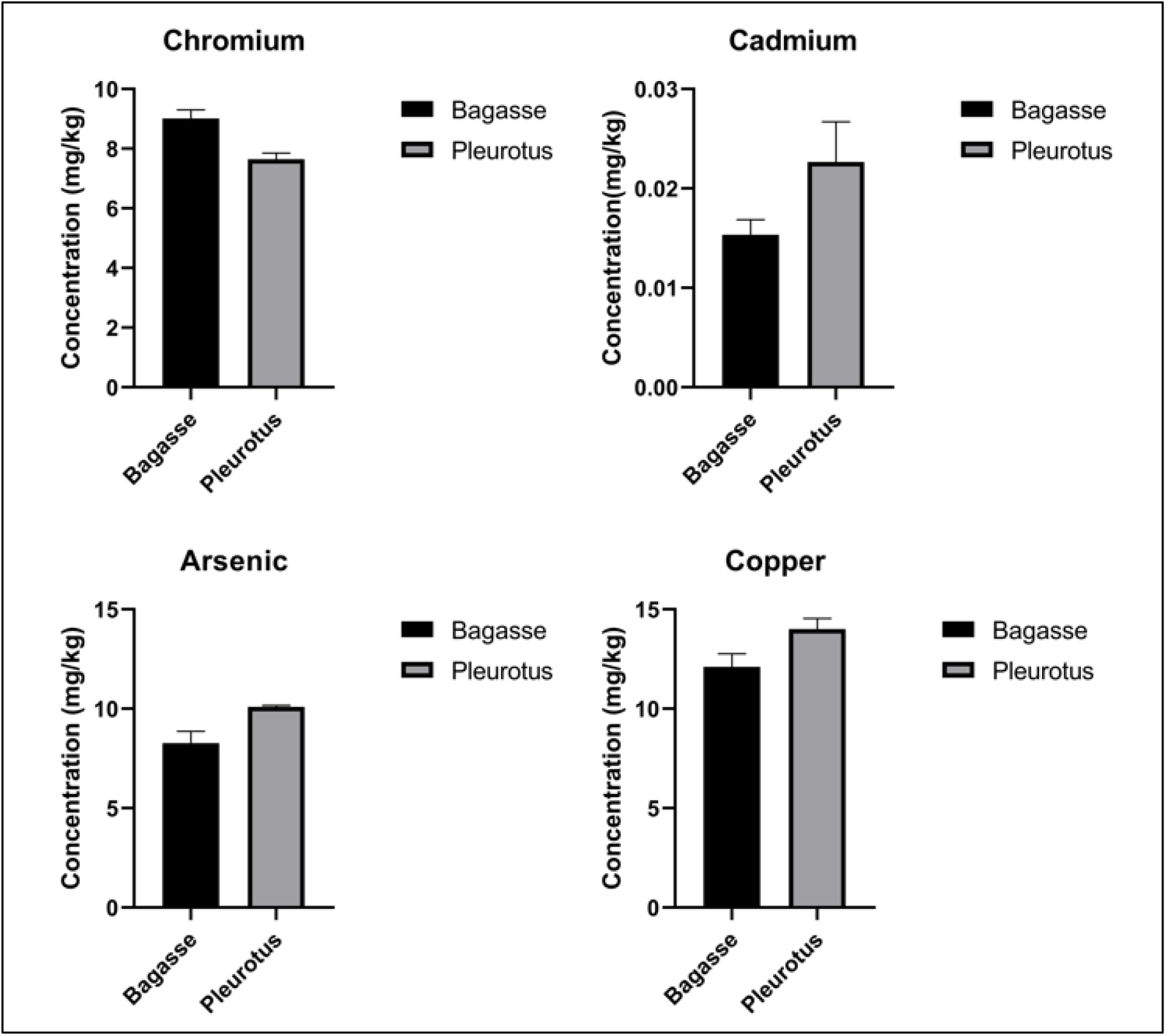
Concentrations of Chromium, Cadmium, Arsenic and Copper in mg/kg dry weight of mushroom and bagasse samples.

As shown in Table I, the metals Arsenic, Cadmium, Copper and Chromium were detected in significant units in both mushroom and bagasse. Mercury was present in insignificant amounts in both the samples.

## 5. Discussions

*Pleurotus ostreatus* is one of the most consumed mushrooms amongst the South East Asian countries mainly China, Japan, and other countries (Raman et al., 2021). Thus, the quality control of such a crop holds immense importance. Bagasse has been used as a substrate for mushroom growth since a long time and is well suited for mushroom growth due to its high nutritional content. Study shows dry bagasse consists of 45% cellulose, 28% hemicellulose, 20% lignin, 5% sugar, 1% minerals, and 2% ash (Mahmud & Anannya, 2021). Thus, any day it is more commercially viable option than conventional substrate straw. Our study also replicates the same fact, where the mushrooms achieved their full size within 25 days compared to the conventional time period of 2 months. Though the increased growth rate proves bagasse to be a promising substrate, given the increased demand for food supply. Thus, it is important to check the metal ion uptake from the bagasse by the pleurotus mushrooms, which is one of the most neglected factors but leaves long term effects in the human body.

In this study, Atomic Absorption Spectroscopy was employed to detect the metal ion concentration in both the bagasse and the mushrooms. The concentration of the toxic metals Cadmium, Arsenic, Chromium, Mercury and Copper were checked for. The Bio-accumulation Factor (BAF) was calculated using the formula (Wang, 2016)-

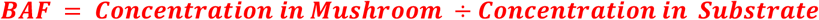

and these metals showed a visible bio-accumulation from the bagasse substrate to the mushrooms. The concentration of cadmium in mushroom was 0.022mg/kg, which was well below the maximum permissible limit of 0.05 mg/kg of dry weight (WHO^1^,1995). The recommended levels for presence of arsenic in mushroom samples is 1.0 mg/kg of dry weight (FSANZ^2^). The concentration of arsenic in mushroom is 10.12 mg/kg which is more than 10 times the maximum permissible concentration and deductive of high arsenic contamination. Copper showed a concentration of 14.29 mg/kg of dry weight which is almost 1.5 times the maximum permissible concentration of 10.0mg/kg (WHO^1^,1995). The recommended levels for presence of chromium in mushroom samples are 0.17 mg/kg of dry weight (WHO^1^,1995). The concentration of chromium in mushroom is 7.59 mg/kg which is more than 40 times the maximum permissible concentration. However, the mercury showed no significant hits, and was thus rendered to be absent in both the substrate and the mushrooms. Studies have established sugarcane bagasse as absorbent for metal ions. The maximum monolayer adsorption capacities of Pb (II) and Ni (II) is found to be 1.61 mg/g and 1.46 mg/g respectively in optimum condition. In another study the concentration of Cu and Ni (II) is in the range of 13.661 mg/kg and 1.56 mg/g respectively (Van Tran et al., 2017). In accordance, this study establishes the presence of metal ion toxicity in sugarcane bagasse used to grow mushrooms.

The bioaccumulation factor calculated for individual metals are of utmost interest. In accordance with the previous studies the bioaccumulation factors were calculated. Biomagnification of metal ions can be very harmful, upon entering the feeder pathway they accumulate in a multiplicative manner and causes various disease like Itai itai (Cd), Arsenicosis (As), Minamata (Hg) etc (Balali-Mood, 2021). Upon assessing the Bio-accumulation factor, which should ideally be ≼1, the BAF in different metals was found to be 1.48(Cadmium), 1.22(Arsenic), 1.16(Copper) and 0.85(Chromium). Given the Bio-accumulation factor and metal ion concentration of Arsenic and Copper, it can be already deemed to be toxic, since this will bio-magnify further down the food chain and along with chromium, although it does not have any bio-accumulation, the metal ion concentration itself is toxic for consumption by human beings. In case of Cadmium, although the concentration in mushroom is well beyond the permissible limit, but this metal had the highest bio-accumulation, and if this trend persists down the food-chain can prove to be alarming.

## 6. Conclusion

Heavy metal toxicity and thereby pollution due to their accumulation is rapidly proliferating, which is a major ecological setback in the present era. These toxic factors have given rise to various biotic and abiotic stresses. In this context, mushrooms growing in the industrial bagasse waste that were collected and tested for certain metals (Cd, As, Cu, Hg and Cr) showed results that were surprising as it showed a progressive biomagnification of the immense metal ions concentration in the mushrooms (Rashid et al., 2018).

This proves that there may be severe impact on the ecosystem as animals consuming such mushrooms having concentrations of heavy metals significantly higher than their individual permissible limits, will be judiciously transferred to the food chain; not to mention human consumption will be equally detrimental. Furthermore, it is well conclusive that these metals are directly coming from the substrate, i.e., bagasse. Some of the scientific methods which could be undertaken to alleviate the high metal ion concentrations from the bagasse, like treating the bagasse with liquid reducing agents (Sodium formaldehyde sulfoxylate), polymer additives (Polyamides and Copolyamides), zinc derivatives (Zinc oxide, Zinc oxide RAC, Zinc Carbonate), nylon additives (Caprolactam), refined silicon alleviation treatment, etc. (Qasem et al., 2021). Understanding of local ecology, risk factors and toxicity in environmental cultivation is essential to initiate a pollution free feeder pathway so that the harmful outbreaks can be prevented.

Understanding of local ecology, risk factors and toxicity in environmental cultivation is essential to initiate a pollution free feeder pathway so that the harmful outbreaks can be prevented.

## Footnotes

1 World Health Organisation

2 Food Standards Australia New Zealand

## Notes

### Competing Interest Statement

The authors have declared no competing interest.

